# Optimal seasonal routines across animals and latitudes

**DOI:** 10.1101/2020.08.02.213595

**Authors:** Uffe H. Thygesen, Andy W. Visser, Irene Heilmann, Ken H. Andersen

## Abstract

Most animals face seasonal fluctuations in food availability and need to develop an annual routine that maximizes their lifetime reproductive success. Two particularly common strategies are reducing energy expenditure and building storage to sustain the animal in meager periods (winters). Here, we pose a simple and generic model for an animal that can decide, at each time during the season, on its level of foraging effort and on building energy stores. Using dynamic optimization, we identify the optimal annual routines that maximize the trade-off between energy and mortality over a life-long horizon. We investigate how the optimal strategies depend on the body size and longevity of the animal, and upon the seasonal variability in the environment. We find that with large fluctuations, the optimal annual routine for small animals is to develop a surviving egg/spore stage rather than to attempt to survive the winter. Medium sized animals invest heavily in reserves to allow long hibernation, while larger animals only need smaller reserves and a shorter hibernation period. In environments with smaller fluctuations, organisms do not need energy stores or hibernation but reduce foraging activities during spring and summer where their fitness is highest. Our optimization model can be used as a null hypothesis to explain the annual routines of animals of all body sizes across the globe.

## 1 Introduction

The seasonal cycle is one of the strongest and most pervasive environmental variations affecting living organisms in nature. The predictability of the seasonal cycle makes it possible for organisms to develop elaborate adaptations to the changing conditions. The direct manifestation of the seasonal cycle is changes in light and temperature, however, the more important forcing for many animals is the changes in resources, be it primary production or other organisms. Typically, the seasonal resource cycle alternates between a feast period (around summer) and famine (winter). The most prominent strategies to deal with the variable resource environment is to make reserves during the end of the feast to survive the famine and to enter a form of hibernation during the famine.

Such annual routines have been described theoretically as an optimization problem (Feró et al., 2008; McNamara and Houston, 2008). The model organism has mainly been small birds, where the annual routines revolve around making fat reserves, optimal moulting time (Barta et al., 2006; McNamara et al., 2008), or hypothermia (lowered body temperature) during winter (Clark and Dukas, 2000; Welton et al., 2002). Few models of annual routines exists for other animals than birds. One exception is copepods, that vary reserve investments and timing of reproduction (Varpe, 2012), or their reproductive mode (Sainmont et al., 2014) according to the seasonal cycle. In both cases (birds and copepods), the examples are environments with strong seasonal fluctuations, either in high latitude temperate systems (birds) or arctic environments (copepods). These cases, however, occupy only a small corner of the variability in life histories and seasonal fluctuations, by describing short-lived animals – between one and a few seasons – living at high latitudes with strong seasonal fluctuations in resource availability.

Development of annual routines are not limited to birds and copepods, but is something that almost all animals do to various degrees. The effect of the seasonal variation is largely influenced by the animals’ life span: for a short-lived animal, the winter is experienced as a very long period of famine spanning a significant part – or the entirety – of the animals’ life, while for a long-lived animal the winter is a recurrent pattern. The effect of the season evidently varies globally with extreme variation at the poles and in continental habitats and lesser in the tropics and in the ocean. Thus organisms with different affected depending on their habitat and their life span.

Here we generalize previous models of optimal annual routines of birds and copepods by considering the effect of seasonal variation on all kinds of animals, from short to long lived, and at different levels of seasonal variation. We use the generic model to form hypotheses about the degree by which seasonal variation affects the life history of the entire animal kingdom on earth. We use optimal foraging theory (Stephens and Krebs, 1987) to predict the optimal decisions of an animal during the season: whether to forage or rest, and whether to build up reserves. The model is formulated generically by using body mass and metabolic theory (Brown et al., 2004) to describe differences in life span, and by considering the environment as periodic variation in resources.

## 2 Optimal allocation to storage and foraging

We consider an adult individual living in a seasonal environment which schedules its foraging effort and its building and use of energy reserves over the season, aiming to maximize its lifetime reproductive success (Fig. 1). Seasonality is expressed through temporal variation in food availability while mortality risk is assumed constant. At each instant, the animal chooses its foraging effort, which determines the amount of energy it has available to maintain metabolism. Next, if there is any surplus energy, the animal chooses whether to allocate it to immediate reproduction or to build up stores to endure periods with energy deficits. We aim to identify how the optimal strategy, i.e. the foraging effort and the allocation of available energy, varies over the season. To this end, we use standard methods from state space modeling (Clark and Mangel, 2000; Houston and McNamara, 1999), where the state at each time is the amount of energy the animal has stored, and we use dynamic programming (Bertsekas, 2005) to solve the optimization problem.

**Figure 1:**
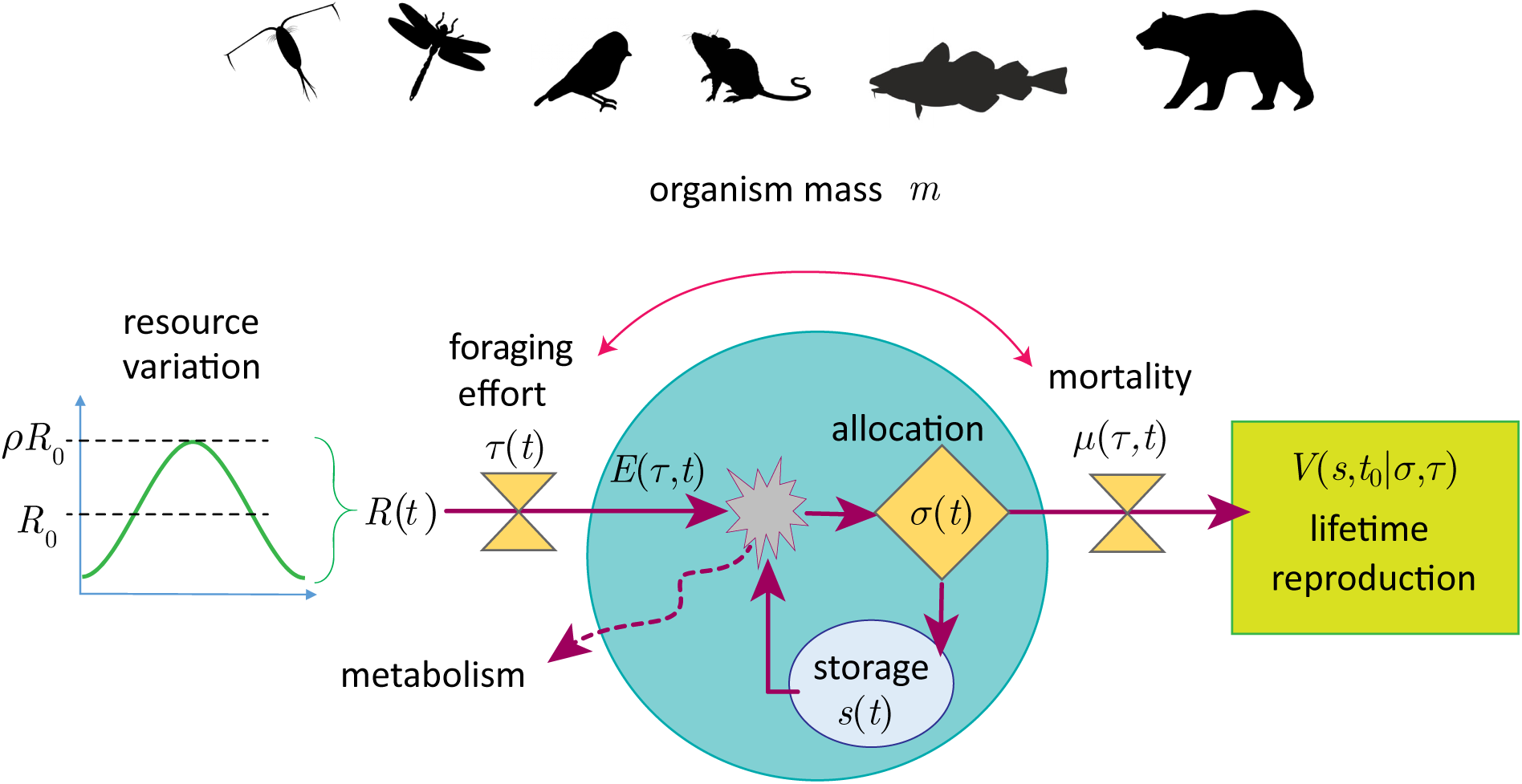
Conceptual sketch of the model. An organism of mass *m* is confronted by seasonally varying resource *R*(*t*). The organism can regulate its foraging effort *τ*(*t*) which effects its energy acquisition but also its exposure to mortality risk. From its energy stream, it pays metabolic costs and can subsequently allocate surplus to storage or immediate reproduction.

Two principal parameters characterize the problem faced by the animal: The relative magnitude of fluctuations in food availability over the season, and the body size (mass) *m* of the animal. We examine how these two parameters effect the optimal strategy. The degree of seasonal fluctuation correlates with latitude, in that polar habitats experience larger fluctuations than lower-latitude habitats, and for a given latitude, oceanic habitats display smaller fluctuations than terrestrial ones. Our model of seasonal fluctuations combines all seasonal patterns in the habitat in a single dimensionless parameter *ρ*, which quantifies the amplitude of fluctuations in food availability measured relative to the average level. The body size of the animal, in turn, defines physiology (consumption rate and metabolism) and predation risk. We consider body size a parameter, not a state variable; i.e. we do not model the growth of the animal. We employ metabolic scaling rules (Brown et al., 2004) to determine how body size affects vital rates. These scaling rules are generic in nature and posit that the speed of metabolic processes (measured in mass per time) scale with body mass *m* roughly as *m*^3/4^. Consequently, rates (units of per time), such as mortality, scales as *m*^−1/4^. It follows that the lifespan scales as *m*^1/4^ – evidently larger organisms have a longer lifespan than smaller organisms. In combination, the magnitude of seasonal fluctuations and the time scale of the animal determine how and to which degree the choices of the animal depend on the time of year.

### 2.1 Energy budgets, reserve dynamics, and the optimization problem

Here, we describe how energy budgets and reserves evolve over the season. We follow state space formalism to describe the animal, with the state *s* ≥ 0 being the energy storage of the animal. As long as the animal is alive the storage evolves according to

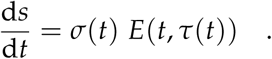

Here, *E*(*t, τ*(*t*)) is the total assimilated energy at time *t*, with *τ*(*t*) being the foraging effort at time *t*, expressed as the fraction of time spent foraging. In turn, *σ*(*t*) denotes the fraction of this energy used for building storage. Both *σ*(*t*) and *τ*(*t*) are decision variables which the animal at each instant chooses from the unit interval [0, 1].

If the assimilated energy is negative, *E*(*t, τ*(*t*)) < 0, we demand *σ*(*t*) = 1, thus forcing the animal to use of its storage. If the assimilated energy is positive, then the animal may choose which fraction *σ* ∈ [0, 1] of this surplus energy that it uses for building storage, while the rest, (1 − *σ*)*E*, goes to instantaneous reproduction. The animal dies if its storage is totally depleted, i.e. *s* = 0. When the animal has positive storage, *s* > 0, its mortality rate is *µ*(*t, τ*(*t*)), i.e., the probability of dying in a short time interval d*t* is *µ* d*t*. Note that the mortality depends on the foraging effort, which leads to an energy-mortality trade-off.

The animal’s goal is to maximize its expected reproductive output during its lifetime by choosing strategies for allocation *σ*(*t*) and foraging effort *τ*(*t*) in each time *t* ≥ *t*_0_, where the initial time is *t* = *t*_0_. The animal dies at a random time *T*, which we integrate out by taking expectation w.r.t. *T*. The expected energy allocated to reproduction for its remaining lifetime is

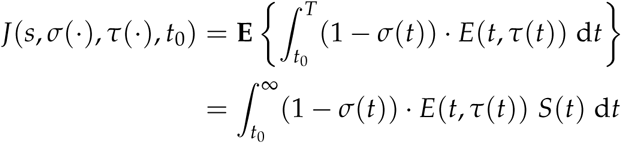

where *s* is the storage at time *t*_0_, **E** denotes the expected value w.r.t. *T*, and *S*(*t*) is the survival function, i.e. the probability of being alive at time *t* ≥ *t*_0_, which can be written 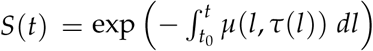.Note that the expected reproductive output *J* depends on the entire functions *σ*(*t*) and *τ*(*t*) for *t* ≥ *t*_0_.

To summarize our problem, we wish to maximize the expected energy allocated to reproduction during the animal’s entire lifetime:

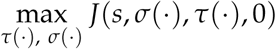

where we assume the animal is born at time *t* = 0. The optimization is subject to conditions for the storage *s*(*t*) described above, and the constraints on the feasible instantaneous decisions, which depends on time:

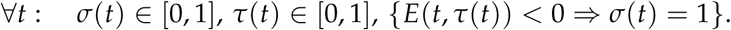

#### Hamilton-Jacobi-Bellman equation

This optimization problem can be solved with the theory of dynamic optimization using the Hamilton-Jacobi-Bellman theorem (Bertsekas, 2005; Clark and Mangel, 2000; Houston and McNamara, 1999). We define the fitness *V* as the expected energy allocated to reproduction in the remaining lifetime of the animal, assuming that the animal behaves optimally, i.e.

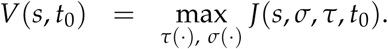

The fitness of a dead animal is zero, while the fitness *V*(*s, t*_0_) of a live animal, *s* > 0, is governed by the Hamilton-Jacobi-Bellman equation:

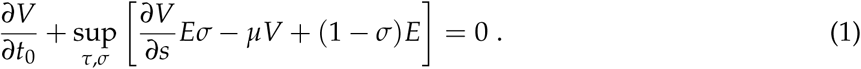

Here, the supremum is over all decisions which are feasible at the instant; in particular, *σ* must equal 1 whenever *E* is negative. The Hamilton-Jacobi-Bellman equation is complemented by the boundary condition

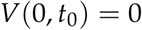

since an animal with storage 0 dies instantaneously and has zero fitness. In the Hamilton-Jacobi-Bellman equation, the term 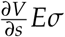 indicates gain or loss of fitness due to building or using storage, −*µV* indicates expected loss of fitness associated with dying, and (1 − *σ*)*E* is the immediate pay-off, i.e. energy allocated to spawning. The Hamilton-Jacobi-Bellman equation expresses a trade-off between sowing and reaping, i.e. building storage to facilitate future spawning or spawning now, and finds from this trade-off the optimal immediate action *τ*(*t*), *σ*(*t*). We solve the Hamilton-Jacobi-Bellman equation numerically as outlined in appendix C.

#### Model components

The available energy is written in terms of a maximum consumption rate *C* (energy per time) and dimensionless feeding levels *f* which is between 0 and 1:

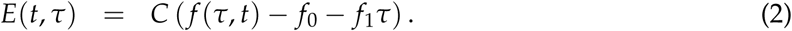

Here, *f* (*τ, t*) is the feeding level at time *t* assuming a foraging effort *τ, f*_0_ defines a critical feeding level where the available energy exactly balances standard metabolism, and *f*_1_*τ* is the extra metabolic cost, expressed as a fraction of *C*, resulting from foraging effort.

For the feeding level *f*, we assume a Holling type II functional response:

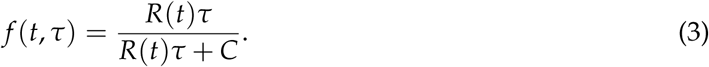

Here *R* is the encountered resource (energy per time) for an animal which forages continuously (*τ* = 1). This varies with the season as:

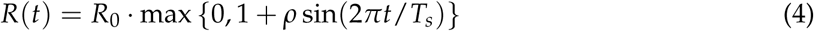

Here *R*_0_ is an average value of the resource, *T*_*s*_ is the period of seasonal fluctuations, i.e. 1 year, and *ρ* is amplitude of resource fluctuations relative to the mean level. The max-function ensures the resource will be non-negative if we increase *ρ* above 1. We use values of *ρ* > 1 to represent environments with prolonged periods without feeding opportunities; i.e., harsh winters.

The mortality *µ* (dimensions: per time) is defined from a background mortality *µ*_0_ and a mortality *µ*_1_ due to foraging effort *τ*

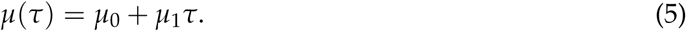

#### Non-dimensionalization and scaling with size

We non-dimensionalize the model using the length of the season *T*_*s*_ as characteristic time and the body mass *m* of the animal as characteristic mass. See details in appendix A. We thus reach the non-dimensional Hamilton-Jacobi-Bellman equation

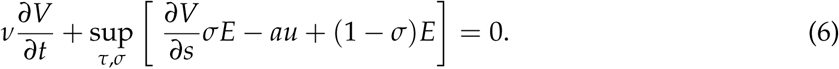

where all quantities are now non-dimensional. The parameter *ν* = *m*/(*T*_*s*_*C*) is non-dimensional and measures the size of the animal relative to the maximum energy consumption during a season and other quantities are non-dimensionalized with *T*_*s*_ and *w*. Also,

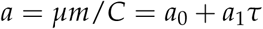

is the non-dimensional physiological mortality, i.e., mortality relative to maximum acquired energy. In a constant environment, a non-dimensional physiological mortality greater than one implies that the expected assimilated energy during the remaining lifetime is less than the body mass of the animal.

We next introduce scalings with size. We write the maximal consumption rate as *C* = *hm*^3/4^. This gives

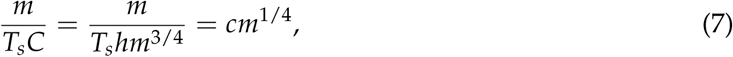

where *c* = 1/(*T*_*s*_*h*) ≈ 0.025 g^−1/4^ is a “pace of life” constant (see Appendix B for parameter values). The factor *m*/(*T*_*s*_*C*) will have magnitude of order 1 when *m* ≈ 2.5 ton, implying the factor will be small for most of the organisms we consider. When the factor *m*/(*T*_*s*_*C*) is small, the fitness function *V* responds quickly to instantaneous conditions, and the optimal strategy depends only weakly on future changes in the environment, whereas it depends strongly on instantaneous conditions. Conversely, when the factor *m*/(*T*_*s*_*C*) is large, the fitness function will only fluctuate weakly over the season, and at each instant the trade-off between energy and mortality is defined by the average fitness *V*.

## 3 Results

The model predicts a wide variety of seasonal strategies (Fig. 2.1). Animals typically have high foraging effort in the spring and fall where the resources are lower. During winters, with strong seasonal fluctuations, the resources are too small to allow the expenses of feeding, and foraging ceases altogether. During the summer, when resources are abundant, the functional response saturates and it pays off to lower foraging effort to reduce mortality. Allocation to reserves occur in the autumn, just before the effort goes to zero. The acquired resources are then used to fuel the winter metabolism in the absence of an energy input from foraging.

Storage and hibernation periods vary with the body mass (Fig. 3). This variation is largely a result of the assumption of decreasing metabolism per body mass for larger organisms where surviving a winter period of fixed length requires less resources per body mass for a large animal than for a smaller one. Therefore storage declines with body mass and hibernation periods become shorter. Small animals subject to strong seasonal fluctuations do not attempt to survive the winter (the top left corners in the in Fig. 3), and therefore do not build storage.

**Figure 2:**
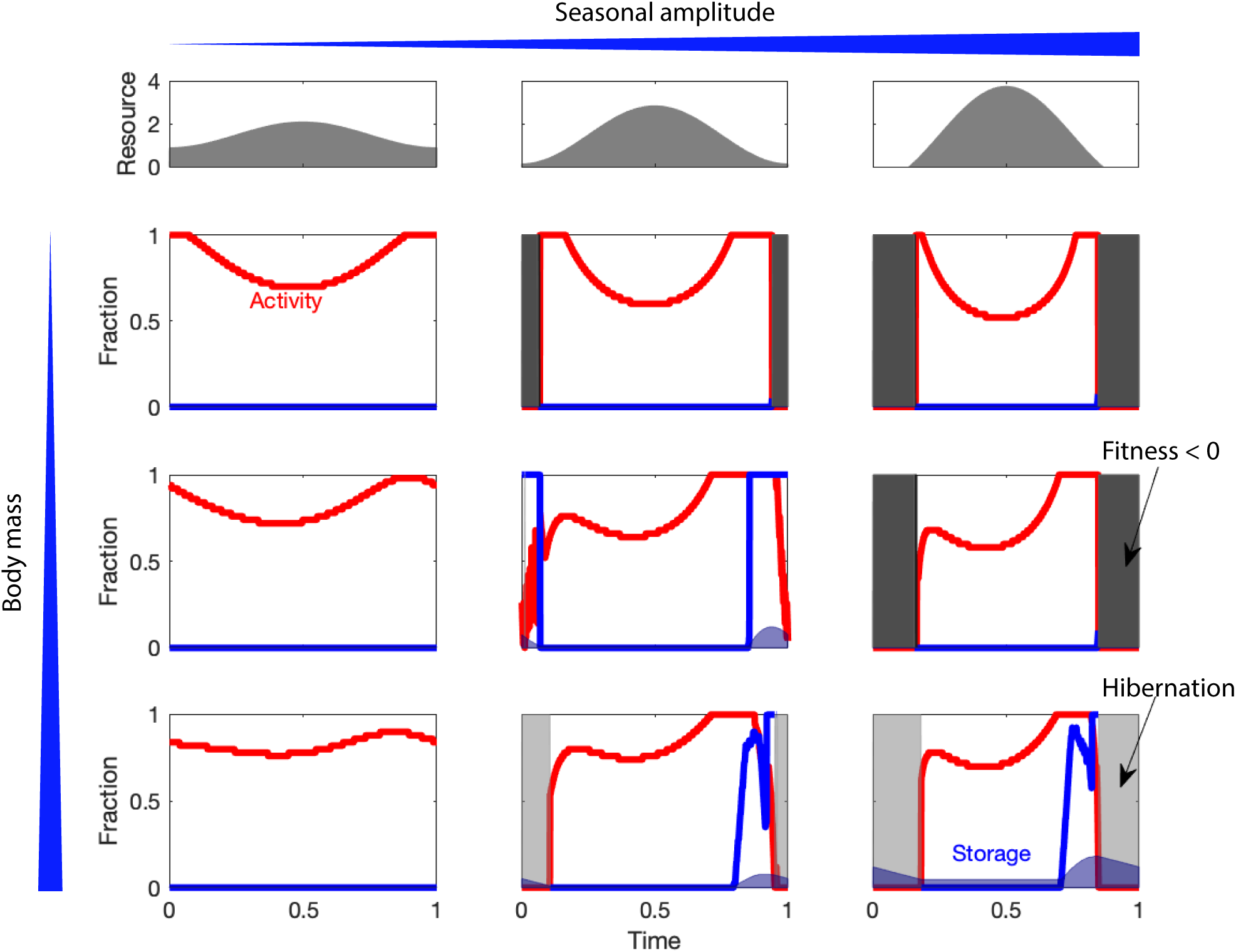
Optimal seasonal routines for animals of different sizes (rows; 0.1, 100, and 10000 g) in increasingly strong seasonal resource environments (columns; *ρ* = 0.4, 0.9, and 1.5). The top row shows the seasonal resource environment. Each panel shows the foraging effort (*τ*, red), allocation to reserves (*σ*, blue) and the size of the reserves as a fraction of body mass (blue patches). Light grey patches indicates time periods where the animals stops foraging entirely and enters hibernation. Dark gray patches indicates time periods where the fitness is less than zero and where the animals either die and leave resting stages behind or migrate away.

**Figure 3:**
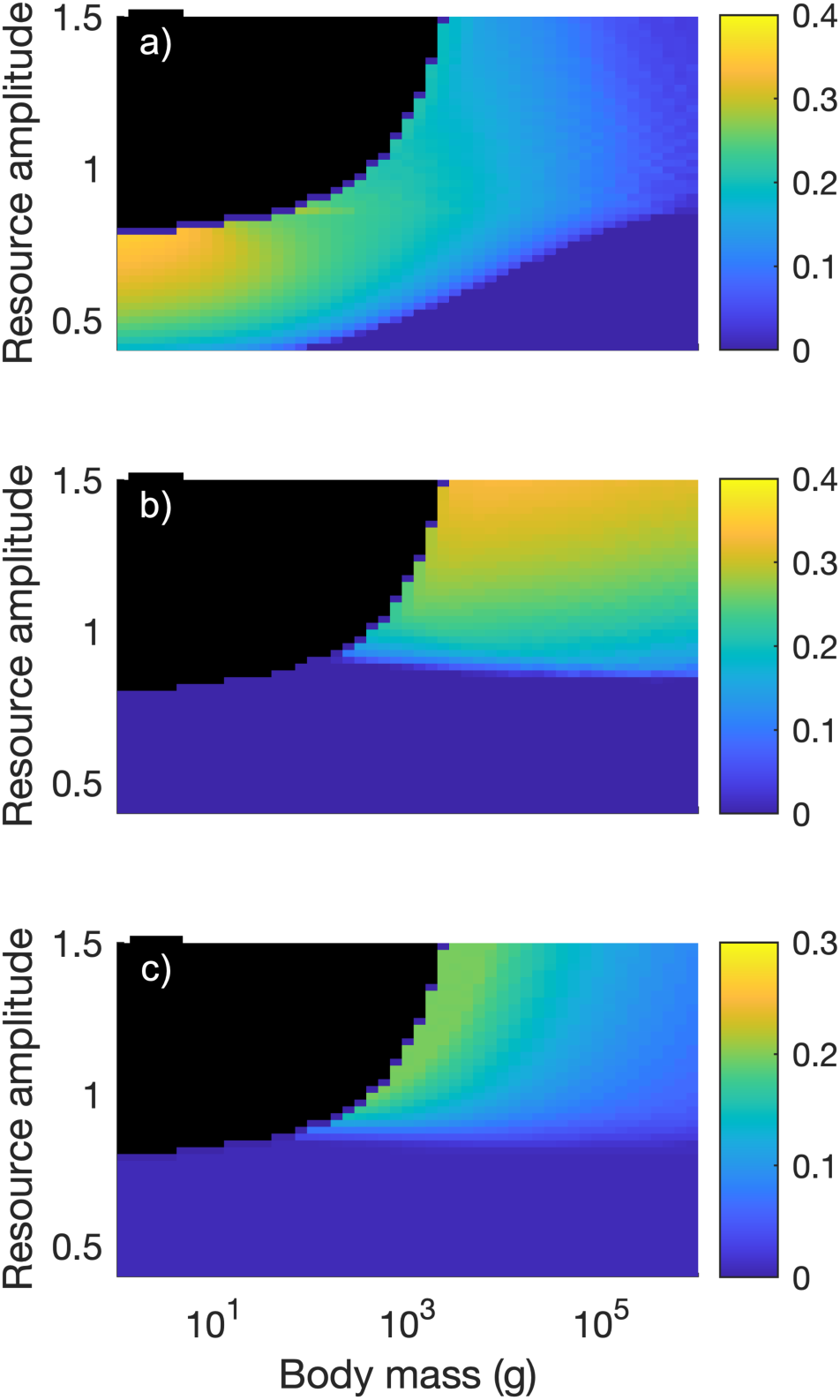
Overview of seasonal strategies for all sizes and seasonal variations. a) The fraction of the season spent foraging at maximum foraging effort (*τ* = 1); b) The maximum size of the storage over the season as a fraction of body mass. c) The length of the hibernation period. The black areas in the top left corners indicate that fitness is less than zero.

## 4 Discussion

From the model results we identify four strategies to deal with a winter famine: 1) increasing the foraging effort to make up for the reduced resources. This strategy occurs in environments with a small seasonal amplitude among organisms of all sizes. Increasing the foraging effort has a cost in terms of increased mortality due to the trade-off between foraging and mortality (5), nevertheless, the increased risk is justified in a life time perspective. 2) Making reserves to fuel winter metabolism. This strategy occurs mainly among larger individuals in stronger seasonal environments, and is accompanied by an increased autumn foraging effort. The prevalence of this strategy among large individuals can be understood by a metabolic argument: surviving a fixed time interval Δ*t* requires a storage that occupies a fraction ∝ Δ*tm*^−1/4^ of body mass. Thus, larger individuals need a smaller fraction of body mass set aside for reserves than smaller individuals, however, the fraction increases proportional to the length of the period the animal has to survive on storage. 3) Stop foraging entirely and go into hibernation. Again, this occurs mostly for smaller individuals and in combination with building storage needed to survive the hibernation. The advantage of hibernation is a reduction in mortality and in metabolism. Hibernation occurs in the model in periods where there are few resources. 4) The last strategy occurs when the fitness drops to zero and the model is unable to find a strategy that leads to survival of the population. This occurs among smaller individuals when they are unable to accumulate sufficient reserves to survive the winter. The four strategies are not distinct but often occur in combination.

The four idealized strategies can be used to interpret the choices made by animals. For example, copepods in high latitudes are larger than in lower latitudes and build bigger reserves (Brun et al., 2016). A particular example is the dominant arctic species *Calanus finmarchicus* and *hyperboreus. C. finmarchicus* builds limited reserves but make a winter hibernation. *C. hyperboreus* is an even larger copepod which make even larger lipid reserves. The smaller *C. finmarchicus* dominates on lower latitudes while the larger *C. hyperboreus* dominates at higher latitudes. The lengthened growth season in arctic environment favours *C. finmarchicus* (Møller and Nielsen, 2019). At very strong seasonal fluctuations even large animals need to enter a hibernation period, such as polar or brown bears.

The last strategy identified by the model (4) indicates that small organisms need to develop alternative strategies to cope with the winter famine. One strategy is for the adult organism to die and leave behind seeds, eggs, or resting spores. Smaller plants and scrubs leave seeds for the next generation even in low seasonal environments. Some copepods species make eggs that sink to depth while the adult perishes (Holm et al., 2018). Many unicellular aquatic protists form resting stages in the form of cysts, e.g., dinoflagellates (Zonneveld et al., 2013) and diatoms (Hallegraeff and Bolch, 1992). Smaller fish species, like capelin (*Mallotus villosus*), have semelparous reproduction where they spawn and die because they are unable to make sufficient lipid reserves to both survive the winter and fuel reproduction. An alternative winter strategy is a latitudinal migration to climates with a less extreme winter. Migration is mainly done by birds but also fish species (e.g. herring, mackerel, and garfish) follow a seasonal peak in primary production across latitudes. Migration requires that the animal is able to make sufficient storage to fuel the migration and it is therefore mainly an option for larger organisms due to the metabolic argument given above. Smaller land-bound organisms, such as rodents, increase their effective reserves by building winter supplies. In this manner their foraging is not limited by their functional response (assuming that it represents gut limitation and not handling limitation) and they can build effective reserves larger than their own body mass. These particular challenges for small and short-lived organisms were also noted by Pianka (1970) associated with *r*-strategies.

Our model is simplistic and aims to describe generic patterns in responses to seasons with a minimal amount of detail. A fruitful avenue of future research could be to make the model specific for a given animal in a given environment, which would complement previous studies that take as starting point specific cases. For the environmental fluctuations, the degree of seasonality varies along a latitudinal gradient, but also between continental, coastal, and oceanic habitats, and our representation of these fluctuations is suitable only at the generic level. Specific studies could detail these fluctuations and include a gonad building between reproductive seasons (Thygesen et al., 2005), or maintain continuous reproduction but let the fitness of the offspring depend on the season. Fluctuations in mortality over the year could also be included, reflecting presence and activity of predators, as well as vulnerability to predation. At the generic level and at least for larger animals, we expect that inclusion of fluctuating mortalities only alters the results quantitatively, in that the optimal strategy would be to reduce foraging effort during periods with increased risk. For smaller animals in strongly seasonal environments, the risk of dying of starvation implies fluctuating food availability leads to different strategies than fluctuating mortalities. For specific animals, fluctuations in mortality is probably necessary to obtain a quantitative match between model predictions and observations.

An even more elaborate model would consider the seasonal game between predators and prey, either in a generic or specific setting, where the the optimal foraging effort of predators depends on the foraging effort of prey and vice versa. An other extension would be to include growth and life history, considering structural body mass an evolving state rather than a fixed parameter. This would increase the fidelity of the model, in particular for short-lived animals.

At a technical level, our model operates in continuous time and continuous state space, while the majority of similar studies have followed Clark and Mangel (2000) and used a discrete setting. The two formulations are, of course, essentially analogous and the difference boils down to whether time steps and number of storage levels are consider numerical parameters or model parameters. We find that the continuous formulation present somewhat cleaner conceptual framework, which is particularly appealing when operating at the generic level.

## 5 Conclusion

We have presented a minimalistic model which explains generic patterns in how animals can and should cope with fluctuating environments. We have found that the optimal responses can, qualitatively, be categorized into four broad classes, which each are feasible and favorable for animals of different sizes in different degrees of fluctuations: When seasonal fluctuations are modest, foraging efforts decrease during periods with abundant resources, following the varying trade-off between foraging and mortality. Larger animals in strongly fluctuating environments build storages, whereas smaller animals are more prone to going into hibernation. Finally, smaller animals subject to strong fluctuations do not attempt to survive the famine but concentrate on reproducing as long as possible. The model results conform to and formalizes our general understanding of how seasonal fluctuations affect animals, and the model may serve as a useful null hypothesis to interpret observed seasonal strategies of plants and animals globally.

**Table 1:** Parameters in the model. Justification of values in Appendix B

| Parameter | Value |
| --- | --- |
| Standard critical feeding level | $f_0 = 0.1$ |
| Activity critical feeding level | $f_\tau = 0.1$ |
| Basal physiological mortality | $a_0 = 0.3$ |
| Activity physiological mortality | $a_1 = 0.3$ |
| Average scaled resource level | $\tilde{R}_0 = 1.5$ |
| Pace of life constant | $c = 0.025 \text{ g}^{-1/4}$ |
| Seasonal variation in resource | $\rho = \text{free}$ |
| Body size | $m \text{ free}$ |

## Acknowledgments

[Omitted at this stage due to double-blind review]

## A Non-dimensionalization of the model

To non-dimensionalize the model we scale time with the season to obtain non-dimensional time 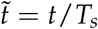, and reserves with body weight to get the non-dimensional reserves 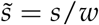.

We describe mortality in terms of the dimensionless “physiological mortality” *a*, using the maximum consumption rate *C* and body weight *w*:

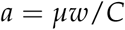

and using this in the equation for *µ*(*τ*) gives

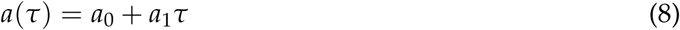

where *a*_0_ = *µ*_0_*w*/*C* and *a*_1_ = *µ*_1_*w*/*C* are background and foraging-related physiological mortalities, respectively.

In the same manner, the encountered resource can be written in non-dimensional form by scaling with the constant *C* giving a non-dimensional level of the food resource 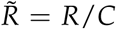. This gives

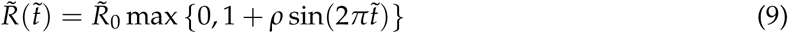

where 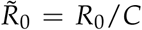 is the dimensionless average resource level. We use 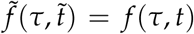 for the functional response as a function of dimensionless time:

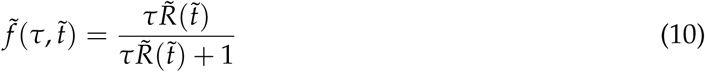

Now the dimensionless energy 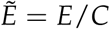 can be expressed as

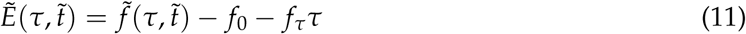

Finally we scale fitness with body weight 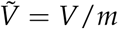. Introducing this in (1) gives:

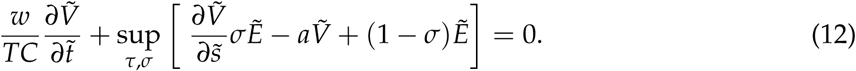

This equation is identical to (1) except for the factor *ν* := *w*/(*TC*), which combines the pace of life *C*/*m* and the length of a season *T*. Aiming for a less cluttered notation, we drop the tilde’s so that *V, t* and *E* from now on refers to non-dimensional quantities.

## B Parameter values

The model requires a specification of the body mass *m* and the amplitude of the resource *ρ*. Additionally, it requires 6 parameters, the metabolic costs *f*_0_ and *f*_1_, the physiological mortalities *a*_0_ and *a*_1_, the scaled resource level 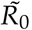, and the pace of life constant *c*. The metabolic costs and the mortalities are non-dimensional constants which are bounded between 0 and 1. The scaled resource is unbounded (but positive), and the pace of life constant is the only dimensional parameter. We use general arguments to find reasonable values of all parameters.

We first consider an animal that forages at maximum rate, i.e., *τ* = 1. For this case we assume that the total metabolism and mortality are split evenly between basal rates and active rates. With this assumption, *f*_0_ = *f*_1_ and *a*_0_ = *a*_1_. We further assume that the total metabolism is on the order of 20% of the maximum consumption rate. This yields *f*_0_ = *f*_1_ = 0.1.

We can consider the level of the resource 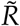 by again considering an animal that feeds all the time (*τ* = 1). Its feeding level *f* ^∗^ is (10):

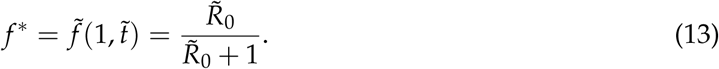

This yields the food consumed relative to the maximum consumption for an individual that forages at maximum rate. This value *f* ^∗^ should, on average, be larger than *f*_0_ + *f*_1_ = 0.2 to ensure sufficient intake to cover metabolism, while still less than the the upper bound *f* ^∗^ ≤ 1. A value of 0.6 seems appropriate – in this case the organism is neither starving nor satiated. This assumption provides a reasonable value for the scaled average resource concentration:

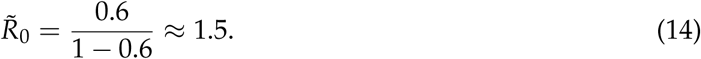

Larger values of 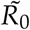 will lead to satiated individuals without the need for seasonal strategies, while smaller values will lead to starving individuals and more extreme seasonal strategies.

To estimate a reasonable value for the overall physiological mortality *a*, we consider first the Darwinian fitness, i.e., the expected number of offspring which will reach maturity:

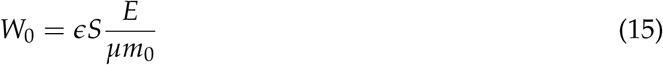

Here, *S* is the survival to maturation, 1/*µ* is the expected remaining lifetime of an adult, so that *E*/*µ* is the expected future reproductive output of this adult. *ϵ* is the reproductive efficiency and *m*_0_ is the mass of an offspring, so *ϵE*/(*µm*_0_) is the expected number of offspring the adult will produce in its remaining life. Considering again an individual feeding constantly and with feeding level *f* ^∗^, we get *E* = (*f* ^∗^ − *f*_0_ − *f*_1_)*hm*^3/4^ (2) and from (7) we have *µ* = *ahm*^−1/4^. Assuming that the physiological mortality *a* is constant during the growth phase of the animal, the probability of survival from size *m*_0_ to size *m* is *S* = (*m*_0_/*m*)^*a*^ (Andersen, 2019). Inserting in (15) we get:

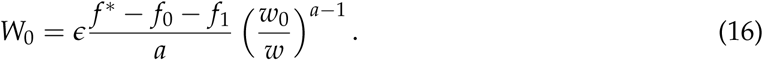

If the population is in steady state, we require that *W*_0_ = 1 and use this to determine *a*. With *ϵ* = 0.2 and *m*/*m*_0_ = 100 (Neuheimer et al., 2015) we find *a* ≈ 0.6 which compares well with empirical measurements on fish and elasmobranchs (Andersen, 2019, Ch. 9). We then get *a*_0_ = *a*_1_ = 0.3.

## C Numerical methods

We solve the Hamilton-Jacobi-Bellman equation numerically using simple and conservative methods; from the point of view of computational methods for partial differential equations, the problem is modest in complexity and computational requirements. The state space is truncated so that the dimensionless storage is bounded above by 0.2. The state space is discretized into 30 grid cells. We time-step the Hamilton-Jacobi-Equation by the explicit Euler method, using a time step of 10^−3^ years. At each time step, and for each possible state, we find the optimal strategy *τ, σ* by brute-force evaluation over a discretized decision space: *τ* is allowed to vary over 51 different values between 0 and 1, while we for *σ* exploit that the objective function is linear in *σ*, so that the maximum is attained for *σ* = 0 and *σ* = 1. The time marching continues for 10 years to ensure that the periodic asymptote is reached. Sensitivity analysis indicates that the results do not depend significantly on numerical choices, i.e. grid sizes, time steps, etc.

